# Continuous monitoring of cerebrovascular autoregulation using functional ultrasound imaging in the piglet brain

**DOI:** 10.1101/2025.09.09.675065

**Authors:** Sofie Dietvorst, Clément Brunner, Dries Kil, Elle Scheijen, Gabriel Montaldo, Bart Depreitere, Alan Urban

**Affiliations:** Department of Neurosurgery, University Hospitals Leuven, Leuven, Belgium; Laboratory of Experimental Neurosurgery and Neuroanatomy, KU Leuven, Belgium; Neuro-Electronics Research Flanders, Leuven, Belgium; VIB, Leuven, Belgium; Imec, Leuven, Belgium; Department of Neuroscience, Faculty of Medicine, KU Leuven, Leuven, Belgium

**Keywords:** Cerebrovascular autoregulation, Real-time brain monitoring, Functional ultrasound imaging, Intra-cranial pressure, Cerebral perfusion pressure

## Abstract

Continuous real-time assessment of cerebral blood flow (CBF) and cerebrovascular autoregulation (CA) remains a major unmet clinical need in acute brain injury. Methods such as laser Doppler flowmetry (LDF), transcranial Doppler, or indirect indices lack accuracy and robustness. Functional ultrasound (fUS) is an emerging modality combining high spatiotemporal resolution, large field-of-view, and sensitivity to blood velocity and volume, making it a promising neuromonitoring tool. Piglets were equipped with arterial blood pressure (ABP), intracranial pressure (ICP), and LDF probes, plus cranial windows for fUS and red blood cell (RBC) flux imaging. CA was challenged by non-pharmacological ABP manipulation via intraaortic or intracaval balloon inflation. fUS hemodynamic parameters were compared with other modaliters across a CPP range of 10–150 mmHg. fUS provided continuous, stable intensity- and velocity-derived parameters across vessels types. CBF estimates correlated strongly with RBC flux and showed reproducibility comparable to LDF, with lower inter-animal variability. Autoregulation breakpoints were reliably identified by fUS, particularly the lower limit, while the upper limit was more variable. Parcellation confirmed robustness of fUS across brain regions. fUS images CBF and CA with higher stability and reproducibility than standard approaches, supporting its applicability for bedside neuromonitoring and clinical translation.

## Introduction

Cerebrovascular autoregulation (CA) is an important physiological mechanism for maintaining an adequate cerebral blood flow (CBF) against changes in cerebral perfusion pressure (CPP). Under physiological conditions, CBF is efficiently stabilized by vasodilation when CPP decreases, and by vasoconstriction when CPP increases^1–3^. However, CA can be impaired in acute (e.g. traumatic brain injury, stroke, intracranial hemorrhage)^4,5^ and chronic pathologies (e.g. vascular dementia, Alzheimer’s disease)^6,7^. In addition, the robust assessment of CA impairment provides a clinically relevant outcome, as it allows the assessment of secondary brain injury in a time window that remains a gray area of clinical monitoring^8,9^. Unfortunately, the CA status is difficult to assess and there is no standard procedure to treat patients for a potential disruption of CA.

To assess CA in its physiological state, one needs to quantify CBF during a natural or induced variation of CPP. However, measuring CBF is technically challenging because: i) it must be continuous and real-time to meet clinical needs, ii) it depends on both blood velocity and vessel size, iii) it preferably includes hemodynamic information as downstream as possible iv) the measurement ideally covers a significant portion of the brain (i.e., large field of view) to avoid too local non-generalizable output. Finally, CBF measurement must be minimally invasive and technically feasible to perform in the intensive care unit.

Since this ideal method for measuring CBF is not yet available, several alternative methods have been proposed using direct measurement (e.g., laser doppler flow (LDF)^10,11^, transcranial Doppler^12,13^) or indirect indices calculated from other readily available parameters correlated with CBF (i.e., Pressure reactivity index (PRx)^14^). The detail and comparison of imaging modalities and strategies for addressing CA have been extensively reviewed^15–17^. Nevertheless, no consensus or gold standard approach has been reached on the different methods and metrics for quantifying CA, reflecting the complexity and high uncertainty of construct validity. The lack of standardization in the measurement of CBF is one of the main difficulties in the fundamental and clinical research of CA. Therefore, the development of new alternatives that meet both CBF measurement (i.e., robust, and standardized metrics) and clinical requirements (i.e., low invasiveness with a large field of view; real-time, continuous, and long-term recording) remains critical and urgently needed^9,18^.

In recent years, a method called functional ultrasound (fUS) has been developed to image changes in hemodynamics (CBV, blood velocity) over periods of seconds to hours^19,20^. In typical implementations, this method combines high spatiotemporal resolution (100µm, 200µs), large field of view (1cm^2^), deep tissue penetration (∼1cm), and high sensitivity to detect and quantify blood in small vessels (tens of µm)^20–22^. fUS scans the brain at a high frame rate of 500Hz. The resulting images are a superposition of tissue and blood echoes. High-pass filtering removes the static signal from the brain tissue and extracts the signal from the moving blood^20,23^. The latter signal carries much information, including frequency associated with blood velocity and intensity associated with CBV, which when computed together provides a robust estimator of the CBF^24^. Taken together, these features make fUS a serious candidate for accurate assessment of CA under pathological states.

To evaluate fUS technology in the context of CA monitoring, we imaged the brain of domestic pigs as a striking model due to its similarities in anatomy and physiology compared to the human brain^25–28^. Simultaneous measurements of CBF assessed by fUS, LDF and surface arteriolar red blood cells (RBC) flux by fluorescence were performed in anesthetized piglets during controlled manipulation of ABP, using an endovascular balloon in either the aorta to increase afterload and mimick hypertensive conditions or in the inferior vena cava to decrease preload and mimick hypotensive cases^29,30^.

## Materials and Methods

### Overview of the experiment

The anesthetized piglet is fixed in a stereotaxic frame and two craniotomies are performed; one for each hemisphere of the brain. In the right hemisphere, the cranial window is closed with an optical chamber suitable for tracking labeled red blood cells (RBC) and vessel diameters for calculating RBC flux by fluorescence microscopy. The left hemisphere is covered with an acoustic window to which an ultrasound transducer is attached for continuous fUS imaging. An LDF probe is inserted onto the dura mater through a small hole in the rostral part of the skull and an ICP probe is placed intradurally next to it through a different burr hole (**Figure 1a)**. The CPP was manipulated by gradual inflation of a balloon placed either at the level of the thoracic aorta for the hypertension model or in the inferior vena cava to mimic hypotensive conditions (**Figure 1b)**. This combination allows simultaneous monitoring of CBF with fUS (**Figure 2a)**, LDF and fluorescence microscopy (**Figure 2b)**, together with monitoring of ICP and other physiological parameters. Piglet CA was monitored using these modalities (**Figure 3**) over a range of CPP from 10 to 150 mmHg by mechanical manipulation of ABP.

**Figure 1.**
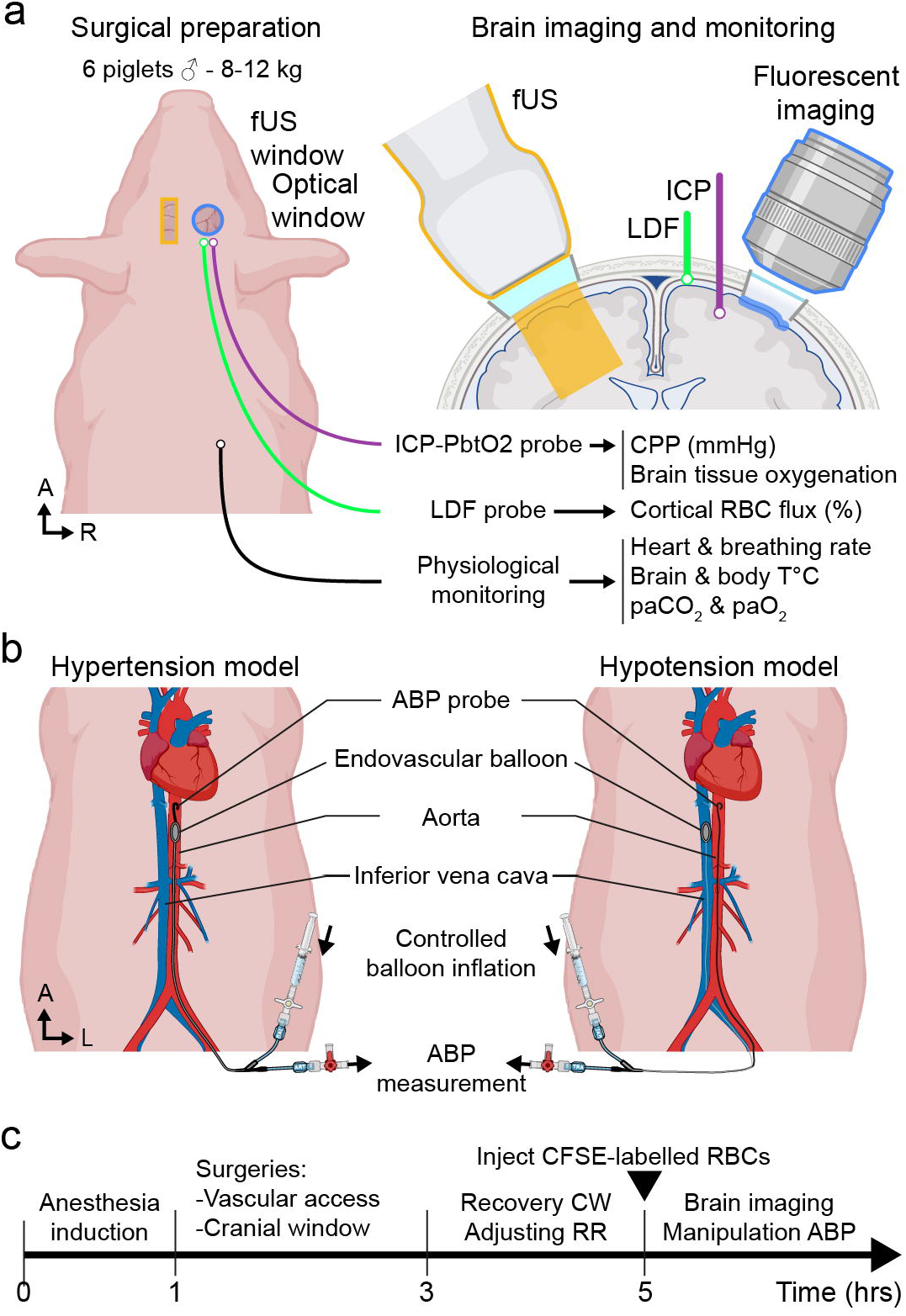
Experimental setup for brain imaging of cerebral autoregulation in pigs. **(a)** *Left;* Illustration of the preparation of anesthetized piglets (i.e., location and size of cranial windows and implanted probes) for multimodal monitoring of brain hemodynamics with joint laser Doppler flowmetry (LDF; green), intracranial pressure and brain tissue oxygen tension (ICP-PbtO_2_; purple), physiological parameters (black), together with functional ultrasound (fUS; orange) and optical imaging (blue). *Right;* Schematic representation of the brain field of view covered by either fUS (orange) or optical imaging (blue). **(b)** Schematic representation (supine view) of experimental models used for controlled manipulation of piglets’ arterial blood pressure (ABP) using an endovascular balloon placed either in the aorta to model hypertension (left), or in the vena cava for hypotension (right). **(b)** Experimental timeline showing the workflow and duration from piglet’s preparation to imaging of pathological hypertension and hypotension conditions. L: Left, R: Right, A: Anterior. All acronyms are detailed in the Materials and Methods section. See **Supplementary Figure 1** for surgical images.

**Figure 2.**
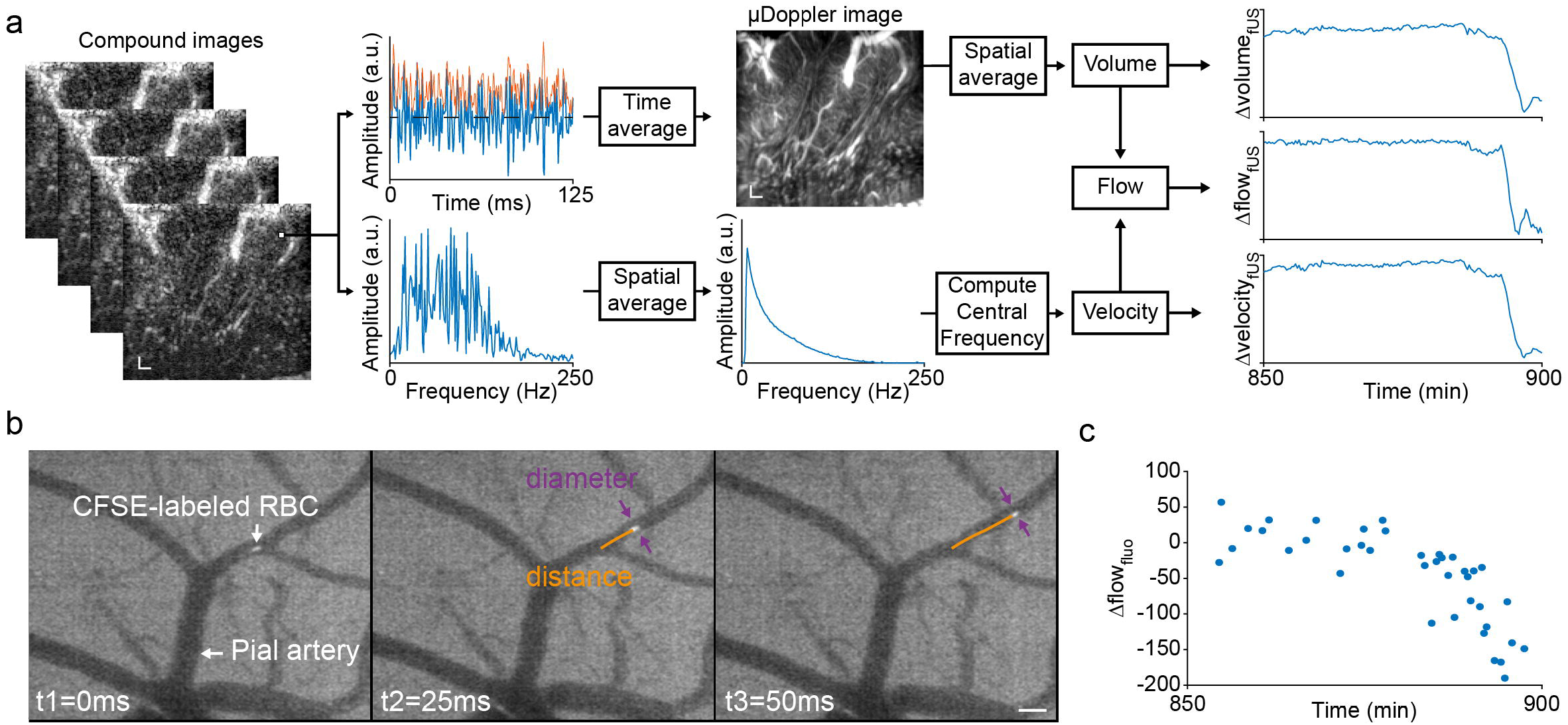
Signal processing of neuroimaging modalities. **(a)** fUS acquisition and data extraction. *Top;* Ultrasound compound images from plane-wave acquisition (see Materials and Methods) are processed with high-pass and singular value decomposition filters and averaged over time to generate a μDoppler image. The temporal change in signal intensity (Δvolume_fUS_ in %) is obtained by spatially averaging successively acquired μDoppler images. *Bottom;* Ultrasound compound images from plane-wave acquisition are decomposed into frequencies by fast Fourier transform to extract the temporal variation in velocity (Δvelocity_fUS_ in %). Changes in signal intensity and velocity are calculated together to generate relative temporal changes in CBF (ΔFlow_fUS_ in %). Scale bar: 1mm. **(b)** Example set of fluorescence images of CFSE-labeled red blood cells acquired in pial arteries during a 50-ms recording period. The distance traveled by CFSE-labeled RBC and the vessel diameter are recorded at 200Hz every 60sec. **(c)** RBC velocity (Δvelocity_fluo_ in %) is calculated and extracted from the distance and diameter measurements along the recording session, including during ABP manipulation. Scale bar: 0.1mm.

**Figure 3.**
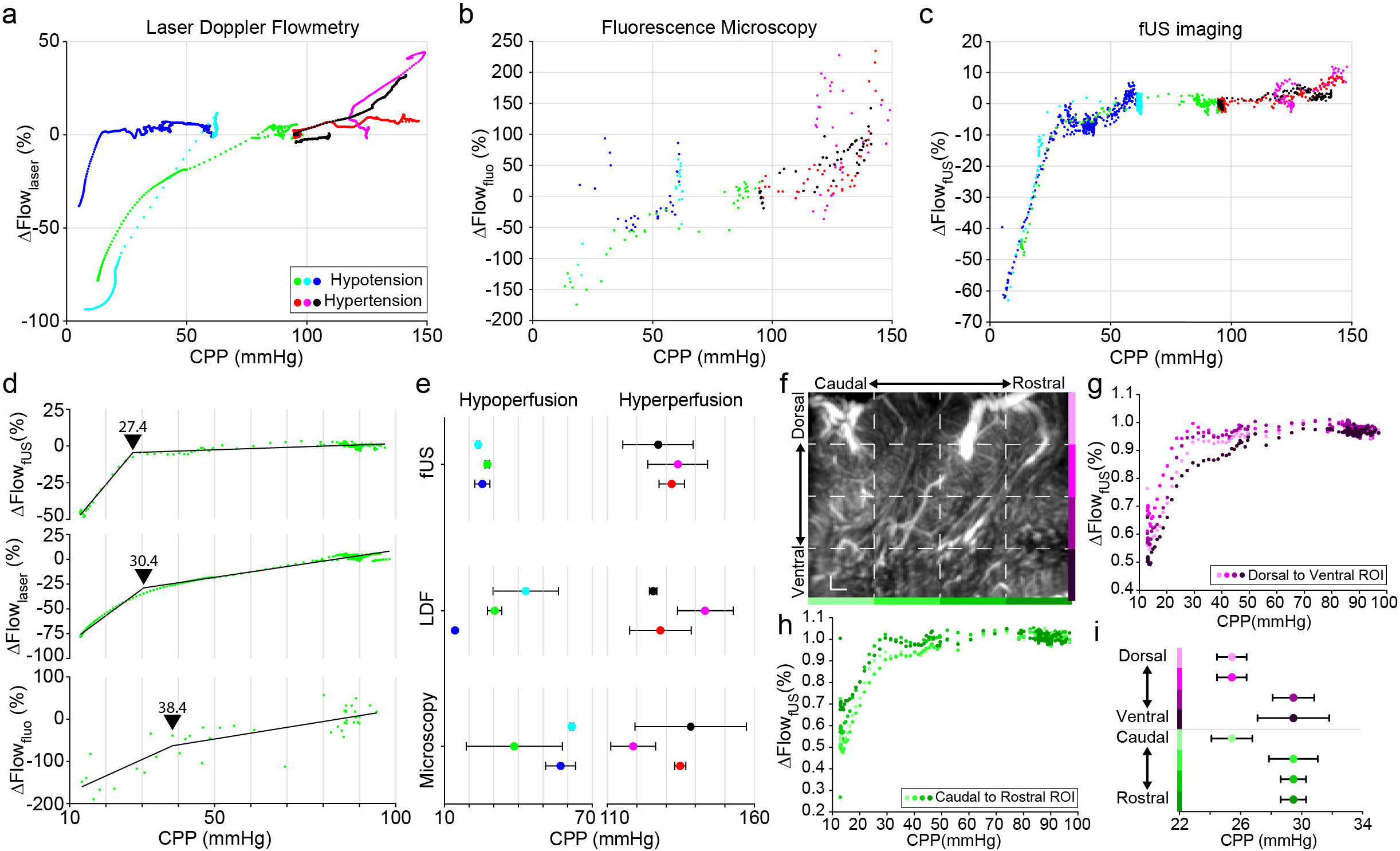
Neuroimaging and monitoring of hyper-or hypotensive pathological conditions. **(a)** Variation of surface CBF measured by LDF (ΔFlow_laser_ in %), **(b)** RBC flow imaged by fluorescence (ΔFlow_fluo_ in %), and **(c)** relative CBF imaged using fUS (ΔFlow_fUS_ in %) measured during change in CPP (mmHg) provoked by the non-pharmacological manipulation of ABP. Data are normalized to the baseline period (pre-ABP manipulation). Color code applies to panels a to c: Blue, cyan, and green dots are measurements from piglets subjected to hypotension (n=3); black, red, and pink are from hypertensive cases (n=3). **(d)** *Top to bottom;* Variation of signal from fUS, LDF and fluorescent microscopy as a function of CPP for one piglet subjected to hypotension (green). Black lines represent the linear regression used for computing the breakpoint (black arrow) and corresponding CCP (mmHg) value for the three methodologies. **(e)** Breakpoint value (mean ± SD) computed for the three methodologies (top to bottom) in each piglet subjected either to hypotension (left) or hypertension (right; same color code applies as in A-C) using Monte Carlo simulation. **(f)** Parcellation of the µDoppler image in dorso-ventral (shades of pink) and rostro-caudal orientation (shades of green) used to extract the signal changes (ΔFlow_fUS_ in %) as a function of CPP in sub-regions of the image, during hypotension. Scale bar: 1 mm **(g-h)** Change in relative CBF (ΔFlow_fUS_ in %) extracted from the **(g)** dorsal to ventral (shades of pink) or **(h)** rostral to caudal (shades of green) regions of interest of the fUS image from the same piglet as in D. **(i)** LLA breakpoint values (mean ± SD) computed from the dorsal to ventral (shades of pink) and rostral to caudal (shades of green) parcellation of µDoppler image using Monte Carlo simulation.

### Animal preparation

#### Ethics

All animal care and procedures were approved by the Ethical Committee for Animal Experimentation at KU Leuven (ECD P107-2019) in accordance with the Belgian Royal Decree (May 29, 2013) and the European Directive 2010/63/EU on the protection of animals used for scientific purposes. All animal procedures were performed under veterinary supervision according to the guidelines of the Ethics Committee.

#### Animals

6-week-old male piglets (domestic swine, Zootechnical Center at the KU Leuven University) weighting between 8 and 12kg.

#### Anesthesia

The animals were prepared as previously described by Klein et al^29,30^. Piglets were premedicated with an intramuscular dose of tiletamine/zolazepam (4mg/kg; Zoletil® VIRBAC, France) and xylazine hydrochloride (1.6mg/kg; Xylazine® V.M.D., Belgium). Vascular access was obtained through an ear vein. All animals were preoxygenated with 100% oxygen. Endotracheal intubation was performed with a cuffed endotracheal tube (Size 4, Mallinckrodt®, Ireland) after a bolus of propofol (2mg/kg; Diprivan® AstraZeneca, UK) and fentanyl (2μg/kg; Janssen-Cilag, Belgium) in prone position. After endotracheal intubation, oxygen was reduced to 40% mixed with air. Anesthesia was maintained by intravenous infusion of propofol (2-4mg/kg/h), midazolam (0.3-0.7mg/kg/h; B. Braun, Germany), and fentanyl (2μg/kg/h - adjusted to response to painful stimuli up to 20μg/kg/h), all by continuous infusion until the end of the experiment. A continuous infusion of pancuronium (0.3mg/kg/hr; Inresa, France) was started before placement of the stereotactic frame and maintained until the end of the experiment. Ventilation was performed with a volume-controlled ventilator (Cato® Dräger, Belgium) delivering tidal volumes of 10-15ml/kg at a respiratory rate of 20-26/min, adjusted to maintain an end-tidal carbon dioxide (CO_2_) tension of 40mmHg, verified by arterial blood gas sampling. At the end of the experiment, the animal was euthanized with an overdose of sodium pentobarbital (150mg/kg; Euthasol® Kela Veterinaria, Belgium). Inhalational anesthetics were not used during any phase of the experiment.

#### Balloon catheter

The left femoral artery was cannulated for placement of an arterial catheter to measure ABP and extract arterial blood gases, the tip was advanced to the level of the thoracic aorta 5cm above the diaphragm. To induce hypertension, the right femoral artery was cannulated for insertion of a 5Fr balloon occlusion catheter (Embolectomy Catheter® 1601-54, LeMaitre Vascular, USA) and advanced into the thoracic aorta at the level of the diaphragm. Similarly, the balloon catheter was inserted into the left femoral vein and advanced into the inferior vena cava at the level of the diaphragm to induce hypotension. A continuous intravenous heparin infusion (50IU/kg/hr) was started at the time of catheter insertion. Animals were then placed in the prone position for stereotactic cranial surgery.

#### Cranial windows

After a midline scalp incision and removal of the periosteum, the cranial bone was exposed. A first craniotomy was placed as previously described^31–33^, anterior to the coronal suture on the right side of the sagittal suture with a high-speed drill continuously cooled with cold saline. Careful hemostasis of the bone edges was performed with bone wax. The dura was removed under optical magnification to avoid damage to the underlying structures. A chamber was placed over the craniotomy and cemented with dental acrylic cement (G-CEM LinkAce®, GC Europe, Belgium). The chamber consists of a stainless-steel ring with 3 injection ports and a central opening of 15mm diameter, which was sealed with a coverslip glass using acrylic adhesive (Histoacryl®, B. Braun, Germany) to prevent CSF leakage (**Figure 1** and **Supplementary Figure 1**). The space under the glass window was filled with artificial cerebrospinal fluid (NaCl 132mM/l, KCl 3.0mM/l, MgCl_2_ 1.5mM/l, CaCl_2_ 1.5mM/l, urea 6.6mM/l, glucose 3.7mM/l, NaHCO_3_ 24. 6mM/l, warmed to 37°C, and equilibrated with 6% O_2_ and 6% CO_2_ in N_2_ to pH 7.35-7.45, pCO_2_ 40-42mmHg, and pO_2_ 42-50mmHg) through the injection ports before sealing with caps. Posterior to the imaging chamber, two 2-mm burr holes were drilled to allow for i) the placement of the intraparenchymal probe for monitoring ICP-PbtO2 (NEUROVENT-PTO, Raumedic AG, Germany) and ii) the 1-mm diameter LDF probe (Moor VMS-LDF1 with VP14-CBF probe, Moor Instruments, Devon, UK) in contact with the dura, avoiding dural and pial vessels (**Figure 1a** and **Supplementary Figure 1**).

A second cranial window was performed on the left hemisphere for fUS imaging. A ∼7×25-mm^2^ bony window was made and the dura was kept intact underneath. A custom 3D-printed transducer holder (**Figure 1a, Supplementary Figure 1**, and **Supplementary Material**) was cemented to the skull. The dura was covered with a layer of 1.5% agarose (Sigma-Aldrich, USA) and a layer of ultrasound transmission gel (Aquasonic Clear®, Parker Laboratories Inc, USA) to ensure acoustic coupling between the tissue and the ultrasound transducer. Finally, the ultrasound transducer was inserted and fixed with screws into the transducer-holder (**Figure 1a)**. Piglets were allowed to recover for 2hr after this procedure.

### Non-pharmacological manipulation of Arterial Blood Pressure (ABP)

ABP was manipulated in two sets of experiments, a hypertensive (n=3) and a hypotensive group (n=3). As previously described^29,30^, the balloon catheter was progressively inflated over a period of 2-3hr by infusion of saline at a rate of 0.1 to 0.3ml/hr using an automated syringe pump and adjusted to the ABP response. The maximum inflated balloon diameter was 13mm after infusion of 1.5ml saline. In the hypertensive group, a progressive increase in ABP was induced by gradual inflation of the balloon catheter in the abdominal aorta, thereby increasing the afterload of the heart and inducing hypertension. In the hypotensive group, a progressive decrease in ABP was induced by slowly decreasing systemic venous return and thus decreasing preload.

### Physiological monitoring

ABP, ICP, PbtO_2_, brain temperature, LDF and fUS signals were monitored continuously. Heart rate was monitored simultaneously by a 3 lead ECG and by arterial pulse wave analysis. Blood oxygen level was monitored using a pulse oximeter and kept at 99–100% during the entire experiment. Inspired and expired concentrations of CO_2_ and O_2_ were monitored with a gas analyzer (Phillips M1026B, Philips Medical Systems, The Netherlands). Arterial blood gases were sampled for verification of continuously monitored end-tidal CO_2_ (PaCO_2_); pH was kept in normal range 7.35–7.45, PaO_2_ at 200mmHg and paCO_2_ at 38mmHg. Rectal temperature was maintained at 38-39°C by a warming mattress and blankets. Continuously monitored signals were stored using ICM+ software (Cambridge University, Cambridge, United Kingdom). ABP, ICP and LDF signals were sampled at 250Hz. The cerebral perfusion pressure (CPP) was calculated as the difference between ABP and ICP^34^. Physiological parameters of each piglets studied can be found in **Supplementary Material**. The LDF probe gives a relative signal proportional to the CBF around the tip of the probe. This signal is named *flow*_*laser*_ (*t*) to simplify the notation.

### Fluorescent labeling of red blood cells (RBC)

After the cranial window procedure, RBCs were labeled with fluorescent dye as previously described^29,30^. Blood samples of 20ml were collected from the arterial line (estimated 4-7% of total blood volume for animals weighing 8-12kg). The samples were centrifuged at 500g for 10min. The erythrocytes were incubated in 40μM carboxyfluorescein diacetate succinimidyl ester (CFSE; Fisher Scientific, Belgium) diluted in phosphate-buffered saline for 15min at 38°C with periodic inversion to ensure uniform labeling. CFSE is a membrane-permeable esterified cytoplasmic dye commonly used to stain viable cells while maintaining functional properties^29,35^. CFSE has an overlapping excitation and emission spectrum with the auto-fluorescent biomolecules in brain tissue. Selective fluorescence filter illumination produces negative contrast delineation of pial vessels^36^. After incubation, RBCs were centrifuged to remove unbound CFSE dye and washed twice with warm PBS at 38°C. Washed and labeled RBC were resuspended in warm PBS to the original volume of 20ml for slow injection through the arterial line over 10min. The procedure was repeated in two steps to minimize systemic effects. The entire RBC labeling procedure takes approximately 2hr.

### Imaging of CFSE-labeled RBC

RBC signal processing was previously described^29,30^. The cortical pial vessels were imaged with an epifluorescence microscope (SMZ18 with P2-SHR Plan Apo 1×, Nikon, USA), illuminated with a solid-state light engine (SOLA SM2, Lumencor, USA), and captured with a high-speed digital CMOS camera (Orca Flash 4.0 V2, Hamamatsu, Japan) controlled by NIS-Elements software (Nikon, USA). A green filter (P2-EFL GFP-B Filter Cube 470-535nm, Nikon, USA) was used to detect CFSE-labeled RBCs. Images were acquired for 60 sec at 170-200Hz and digitally stored for offline analysis.

### CFSE-labeled RBCs quantification

From the microscope, the diameter of the pial arterioles and the velocity of individually labeled RBCs are measured as *diameter* _*fluo*_ and *velocity* _*fluo*_ respectively. Because velocity depends on position within the vessel, i.e., RBC in the center have higher velocity than those near the vessel wall, the velocity of labeled RBCs is averaged over a 5-second period. Imaging periods with less than 3 RBCs are not considered. Flow in the pial artery is calculated from velocity and diameter as follows:

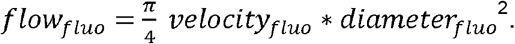

### Functional ultrasound (fUS) imaging

Brain imaging was performed using a linear ultrasound transducer (12MHz, 128-MHz central frequency, 0.125-mm pitch, Imasonic, France) controlled by 128-channel transmit-receive electronics (Vantage, Verasonics, USA). The ultrasound transducer was positioned to image a unique parasagittal brain slice over the left hemisphere (**Figure 1** and **Supplementary Figure 1**) of 16-mm width (antero-posterior axis) and 15-mm depth (dorso-ventral axis).

The fUS sequence was adapted from Brunner, Grillet et al.^22^ The ultrasound scanner imaged the brain at 500 frames/s and the raw images were stored in sets of 250 images (0.5sec acquisition time) for offline analysis. To reduce the amount of data, we saved 2 sets of 250 images per minute for the entire duration of the experiment (∼2hr). Details of the ultrasound sequence parameters can be found in the **Supplementary Material**.

### Hemodynamics parameters extracted from fUS imaging

The ultrasound scanner provides sets of 250 brain images at 500-Hz frame rate (0.5sec acquisition time). These images are a superposition of brain tissue and blood flow. By applying a singular value decomposition filter^23^, we extract the blood signal which is a 3-dimensional matrix *a (x, y, t))* where *x, y* are the spatial dimensions and *t* the time. From this signal, we want to extract a robust parameter to characterize the CA. The two main parameters extracted from the fUS signal are the intensity and frequency of the signal.

The intensity of the signal (also called power Doppler) is an image defined as *I (x, y) = Σ*_*t*_ *a (x, y, t)*^*2*^, which provides an estimate of the CBV and a robust image of the small brain vessels (**Figure 2a)**. A single estimator, proportional to the CBV, is computed as the sum of the intensity of each voxel of the entire image as,

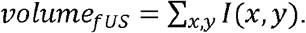

The Doppler velocity is calculated as the center frequency of the Doppler signal (**Figure 2a)**:

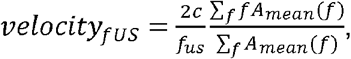

where f is the frequency, c is the speed of sound, and *f*_*us*_ the central frequency of the transducer. *A*_*mean*_ is the averaged spectrum of the Doppler signal over the entire image defined as:

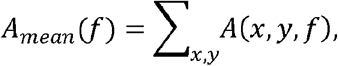

Where *A* is the amplitude of a Fourier’s transform of the Doppler signal *a*.

### Two slopes model and inflection point

In order to fit the curves, a two-slope linear model with an inflection point has been considered. A linear model with a slope *a*_*low*_ was considered below the inflection point *x*_*inf*_ and another linear model In order to fit the curves, a two-slope linear model with an inflection point has been considered. A with slope *a*_*high*_ above the inflection point:

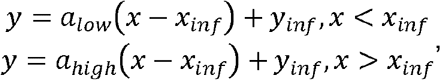

where *x* is the CPP and *y* the flow parameter (from either LDF, fUS, or microscopy). The four parameters *a*_*low*_, *a*_*high*_, *x*_*inf*_, *y*_*inf*_ are fitted by minimal squares. Errors in the parameters are estimated by using a Monte Carlo method.

### Inter-modality comparison and normalization

The three modalities proposed are placed in three different brain locations (**Figure 1a** and **Supplementary Figure 1**) and most of them provide relative measurements. To compare the three modalities, all the measurements were normalized to the baseline in each animal. As an example, the relative variation of the laser Doppler flow at a given time (t) of the experiment is computed as:

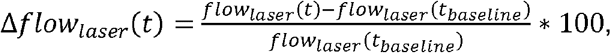

where *t*_*baseline*_ is the baseline period. The aforementioned procedure is then repeated for all parameters derived from all modalities employed. This allows for the expression of all parameters as a percentage of the baseline. **Table 1** provides a comprehensive overview of all parameters that can be compared.

**Table 1.** Parameters measured with each modality.

|  | Laser Doppler<br>Flowmetry | Labeled RBCs<br>imaging | Ultrasound<br>imaging |
| --- | --- | --- | --- |
| Velocity | | $\Delta velocity_{fluo}$ | $\Delta velocity_{fus}$ |
| Volume | | $\Delta volume_{fluo}$ | $\Delta volume_{fus}$ |
| Flow | $\Delta flow_{laser}$ | $\Delta flow_{fluo}$ | $\Delta flow_{fus}$ |
| Diameter | | $\Delta diameter_{fluo}$ | |

## Results

### CBF-CPP relationship and CA breakpoints monitored by fUS, LDF and RBC fluorescence

The proposed methodology enables a continuous recording of the cerebral hemodynamic (parameters are detailed in **Table 1**). For the 6 piglets imaged (n=3 for hypotension and 3 for hypertension), the flow parameters are presented in **Figure 3a-c**, and the velocity, volume and diameter in **Supplementary Figure 2**. As the flow is the only parameter measured by the three modalities, and given its particular importance in cerebral autoregulation, the flow parameter is a primary focus of this work.

The general behavior of the flow in the three modalities (**Figure 3a-c**) follows a triphasic curve as established by Lassen^37^. This triphasic curve consists of a plateau of stable CBF between the upper and lower limits of autoregulation (ULA and LLA, respectively). In the hypotensive experiments, the change in CBF showed a rapid decrease below the LLA breakpoint, whereas in the hypertensive cases, CBF increased more gradually after passing the ULA breakpoint.

In terms of qualitative analysis, the LDF exhibits a stable readout, yet exhibits considerable variation between animals. This variation is likely attributable to the limited field-of-view and the inability to select a specific range of vessels for measurement. Tracking of fluorescent RBCs revealed a considerable variability within the same animal, as evidenced by the reduced number of fluorescent RBCs detected by the method. The ultrasound flow measurement is relatively stable and repeatable due to a combination of a large field of view and high sensitivity.

To quantify the flow curves, both the LLA and ULA breakpoints and their fitting errors were computed using a segmental linear model (see **Materials and Methods**), as illustrated in **Figure 3d**, which depicts the fit for one hypotension case. The breakpoint in this case was easily detected by fUS (27.4mmHg), LDF (30.4 mmHg), and fluorescence microscopy measurements (38.4mmHg).

In the hypotensive model, a comparison of LLA breakpoints between the three modalities (left panel, **Figure 3e**) revealed that fUS exhibited greater consistency and a lower error across animals than LDF and labeled RBCs. In the hypertension model, the comparison of ULA breakpoints (right panel, **Figure 3e**) demonstrated that fUS and LDF measurements exhibited greater equivalence in terms of values and dispersion, and both were superior to the labeled RBCs.

The observed averaged ULA breakpoint exhibited a consistent value across the three methodologies, namely fUS (131.0 ± 3.4 mmHg), LDF (132.3 ± 9.6 mmHg) and RBCs labeling (130.6 ± 10.4 mmHg). However, the averaged LLA is coherent between fUS and LDF, but the labeled RBCs have a higher value (fUS 25.5 ± 1.9 mmHg, LDF at 29.2 ± 14.4 mmHg and RBCs labeling 52.4 ± 12.4 mmHg). It should be noted that the number of animals in each condition is relatively small (n=3), which increases the risk of failing to establish statistical significance. Consequently, the presented results only demonstrate a general tendency for each modality.

### Independence of the image field of view on the fUS signal

The fUS modality measures changes in CBF in a very large field-of-view (∼1cm^2^), especially when compared to LDF and fluorescent microscopy, it captures signals from various type (i.e., arteries and veins) and size of vessels (pial, penetrating and ascending vessels). Therefore, to investigate whether the stability of fUS signals is dependent on image depth and size, fUS images were artificially parcellated into 4 equal horizontal and vertical sections (shades of pink and green, respectively, **Figure 3f**), ΔFlow_fUS_ was measured, and LLA was calculated for each of the parcels. The CBF-CPP relationship was highly similar regardless of the dorso-ventral and rostro-caudal regions of interest (**Figure 3g-h**), as were the LLA breakpoints, which show low variability, with a maximum difference of ∼4mmHg along the dorso-ventral and rostro-caudal parcels of the image (**Figure 3i**). This result supports a high independence of the fUS signal from imaging depth and position within the imaged brain.

## Discussion

In this study, we investigated the feasibility and interest of fUS imaging as a new imaging strategy for continuous assessment of CA status in the context of non-pharmacological hypotension and hypertension in anesthetized piglets. The intensity and spectral signal extracted from fUS imaging were calculated as an estimate of CBF and compared with CBF assessed by LDF and by calculating flux based on microscopic tracking of labeled RCBs and diameter measurement. A segmental linear model was used to evaluate both the CPP values for the lower and upper limits of the CA (LLA and ULA) for which the CA is absent and the CBF-CPP relationship passive^1–3^.

This study was performed with a low number of animals (n=3 for hypotension and 3 for hypertension), therefore it is difficult to draw conclusions about CA physiology itself, which was not the goal of the present study. In a larger study on CA physiology using the fluorescent microscopy method in 20 animals and more arterioles investigated, Klein et al. were able to revisit the Lassen curve and demonstrate differential vasoconstrictive capacity of different arteriolar sizes with increasing CPP and leading to gradual exhaustion of CA capacity on the right side of the CA curve^29,30^. However, the main characteristics of the three modalities can be established from the current analysis. First, CBF, LLA and ULA results obtained with fluorescence microscopy are likely similar to previous recording in similar experimental conditions^30^. Intersubject variability in the limits of CA captured by this means and previously observed was also confirmed in this set of recordings^38^. However, the current fluorescent microscopy settings show more experimental limitations (i.e., RBC labeling and timing, field of view, microscope setting, discontinuous acquisition) with higher instability than LDF and fUS techniques. Such variability can be attributed to the low number of labeled RBCs detected during the experiment (<1 event/sec). In addition, because CBF measurement is highly sensitive to the velocity profile within the imaged vessel, with a maximum velocity in the center of the vessel that decreases to 0 in the wall of the vessel, it is necessary to sample a large number of labeled RBCs randomly distributed in the vessel stream to obtain a stable and thus accurate measure of flow. Second, CBF measurements with LDF show high stability but large inter-individual variability. Indeed, LDF’s limited field-of-view, superficial location, blind and random positioning of the probe is subject to uncertainty when considering the flow measured and may explain the variability of responses among imaged piglets. Finally, fUS is a stable approach to capture CA, providing a robust measure of CBF with accurate detection of the lower limits of CA (i.e., LLA during hypotension) where the inflection is sharp, but reduced performance for its upper limit (i.e., ULA during hypertension) as shown by the larger error intervals. This is consistent with other approaches showing difficulty in deciphering the upper limits of CA^25–28^. In addition, fUS is less subject to inter-animal variability. fUS has a large field of view (1cm^2^) that integrates various vessels (i.e., location, type, size) supporting a stable measure independently of the animal. While the non-pharmacological disruption of CA performed in this work should affect both hemispheres similarly (with no lateralization as observed in stroke or TBI), it is important to note that fUS imaging and other measurements were not conducted in the same hemisphere, which may introduce potential limitations and confounds. Interestingly, our procedure makes each piglet its own control as the multimodal physiological parameters are compared to its own baseline values during the entire experimental duration, before and during ABP manipulation. Taken together, the results support that fUS is an optimal tool for addressing and investigating CA.

We chose a porcine animal model because of its similarities to humans in cardiorespiratory physiology, brain anatomy, cerebrovascular anatomy, and physiology, thus facilitating the clinical translation of such approaches^25–28^. In this work, ABP is manipulated using a mechanical model (i.e., endovascular balloon) to preserve cerebrovascular functions from pharmacological drugs (i.e., catecholamines or vasodilators). Nevertheless, the confounding factor of the anesthetic regimen cannot be excluded but remains close to the protocol used in the neuro-intensive care unit. The experimental procedure has several limitations: first, the placement of the endovascular balloon, which could alter ABP before the start of recordings; second, the surgery and implantation of both open (for fUS) and closed (for fluorescent microscopy) cranial windows, which could locally alter brain physiology. Third, our study was not randomized or blinded due to experimental complexity and constraints, did not mix sexes of animals, and was limited in number (economic constraints), which together could inadvertently introduce experimental bias. In future work, we expect to use a larger number of animals subjected to pathological models (e.g., traumatic brain injury, stroke) to investigate whether the fUS modality can provide an additional benefit in the assessment of CA status in such pathological conditions.

Technically, by reducing the fUS field of view to a quarter of its original size, we confirmed that fUS measurements are not only independent of the rostro-caudal position of the transducer, but also independent of brain depth. In this work, we used an ultrasound transducer with a rather large footprint of 5×12 mm, which may be too large for clinical use. As confirmed by the results obtained after artificial parcellation of the fUS image, the transducer footprint can be reduced to ∼4×6mm. Such a small transducer can contain ∼32 channels, which, combined with a dedicated, simplified, and reduced acquisition system, could be small enough and thus more convenient to be placed and used by clinicians in the ICU.

## Acknowledgements

The authors would like to thank Dr. Stéphanie De Vleeschauwer, veterinarian for the large animal facility of KU Leuven and Inez Feytons, animal caretaker.

## Author contributions

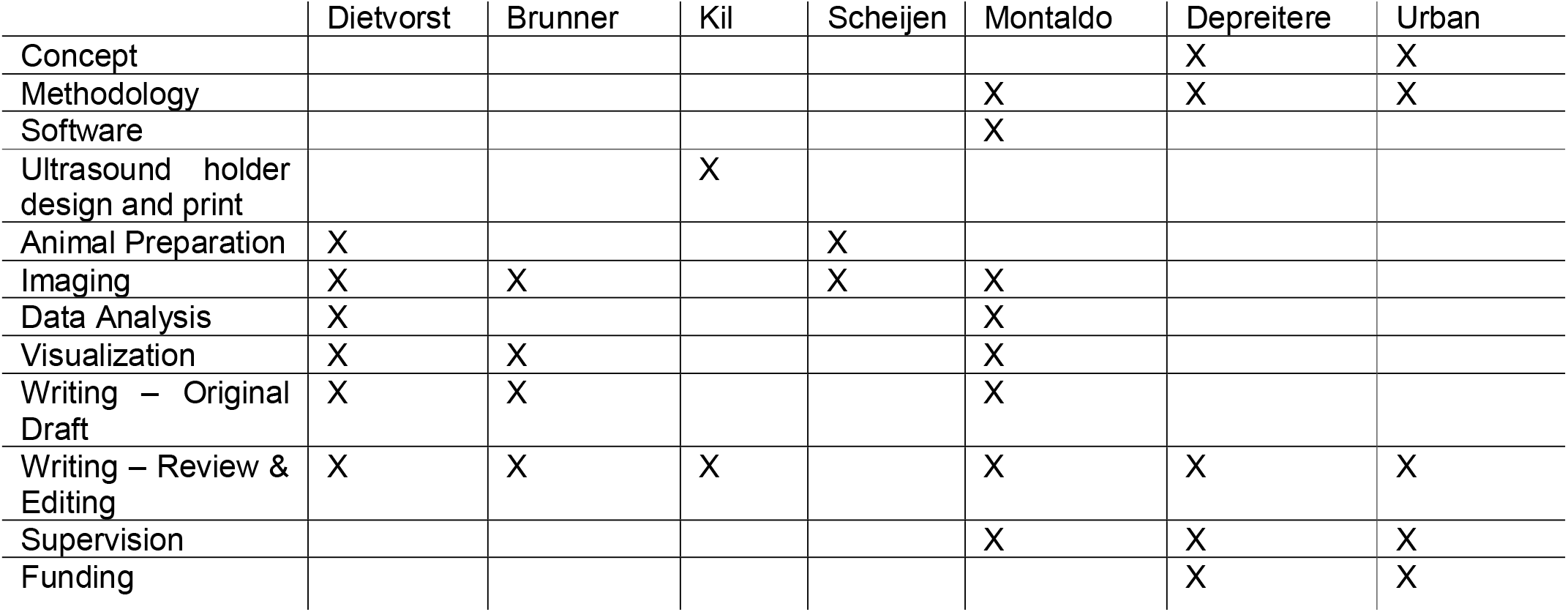

---

**Supplementary Figure 1.** Surgical pictures from multimodal imaging and monitoring of CPP in piglet brain. Left picture shows a top view of the cranial windows (size and location) with holder and chamber surgically implanted. Right picture depicts a back view of the experimental setup on neuroimaging and monitoring including fUS transducer, fluorescence microscope, LDF and ICP-PtO_2_ probe. R: Right, A: Anterior.

**Supplementary Figure 2.** Complementary hemodynamics parameters extracted from fUS (Δvolume_fUS_ and Δvelocity_fUS_ in %; top row) and fluorescence imaging (Δdiameter_fluo_ and Δvelocity_fluo_ in %; bottom row). Color code applies to panels A to C: Blue, cyan, and green dots are measures from pigs subjected to hypotension (n=3). Black, red, and pink are from hypertensive cases (n=3).

## Supplementary Material

1. Computer-aided design files of the transducer holder.
2. Physiological parameters of each individual pigs monitored.
3. Sequence parameters used for fUS imaging.

