## Supplementary figures and images for "Continuous monitoring of cerebrovascular autoregulation using functional ultrasound imaging in the piglet brain"

### 02_SuppMaterial_fUS-holder.jpg

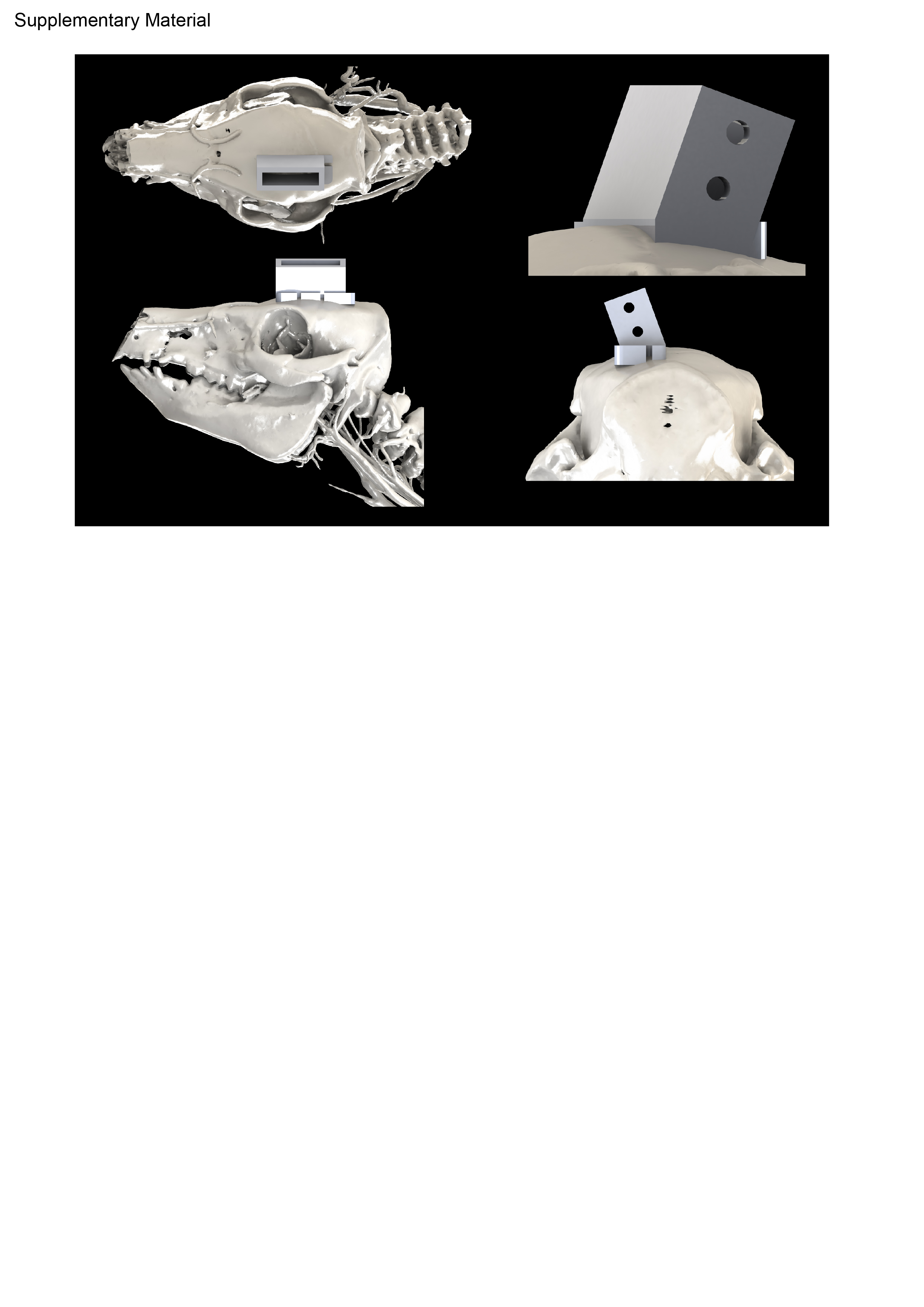

### 03_Pictures_Pig_ProbeHolder_angle.png

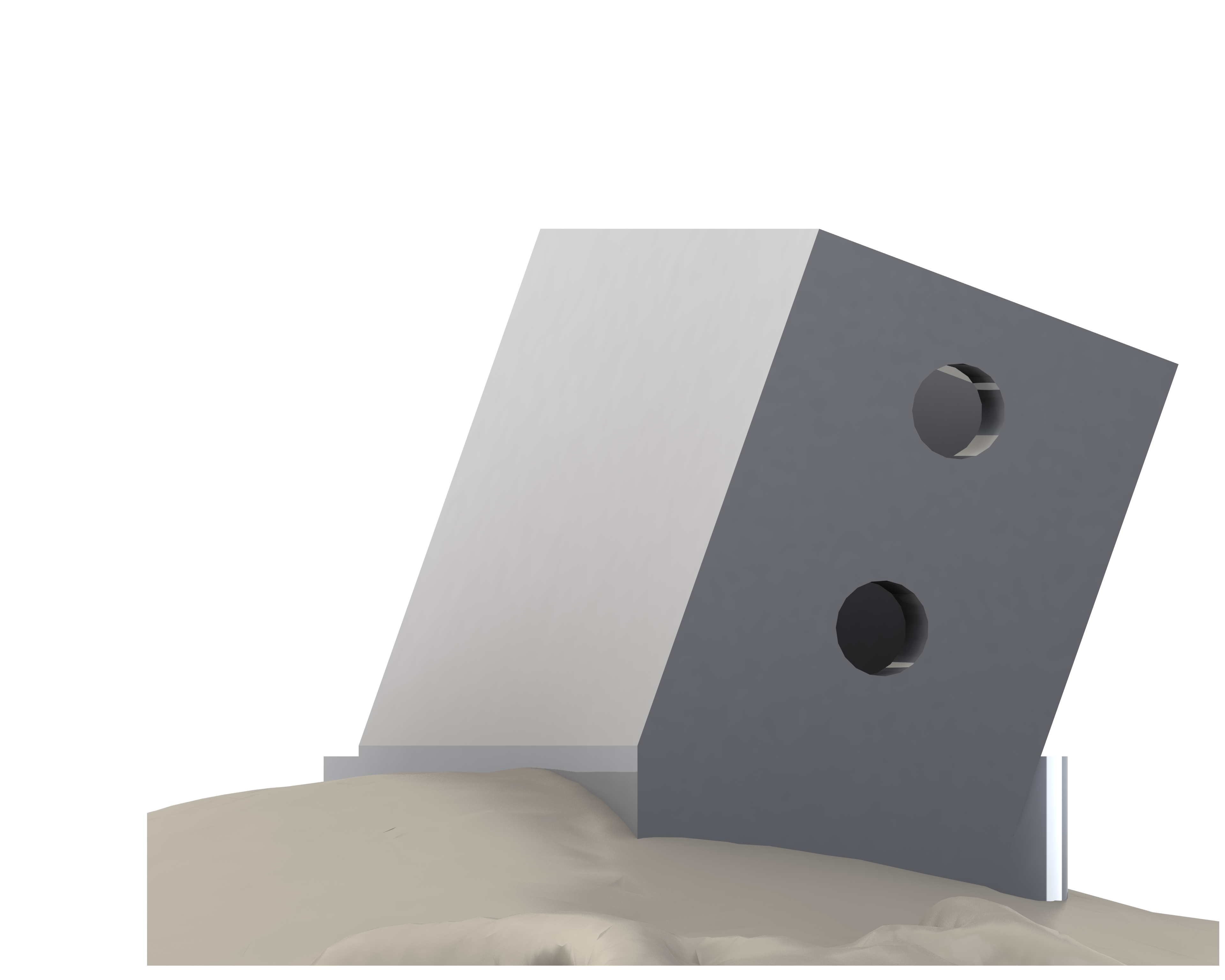

### 03_Pictures_Pig_ProbeHolder_back.png

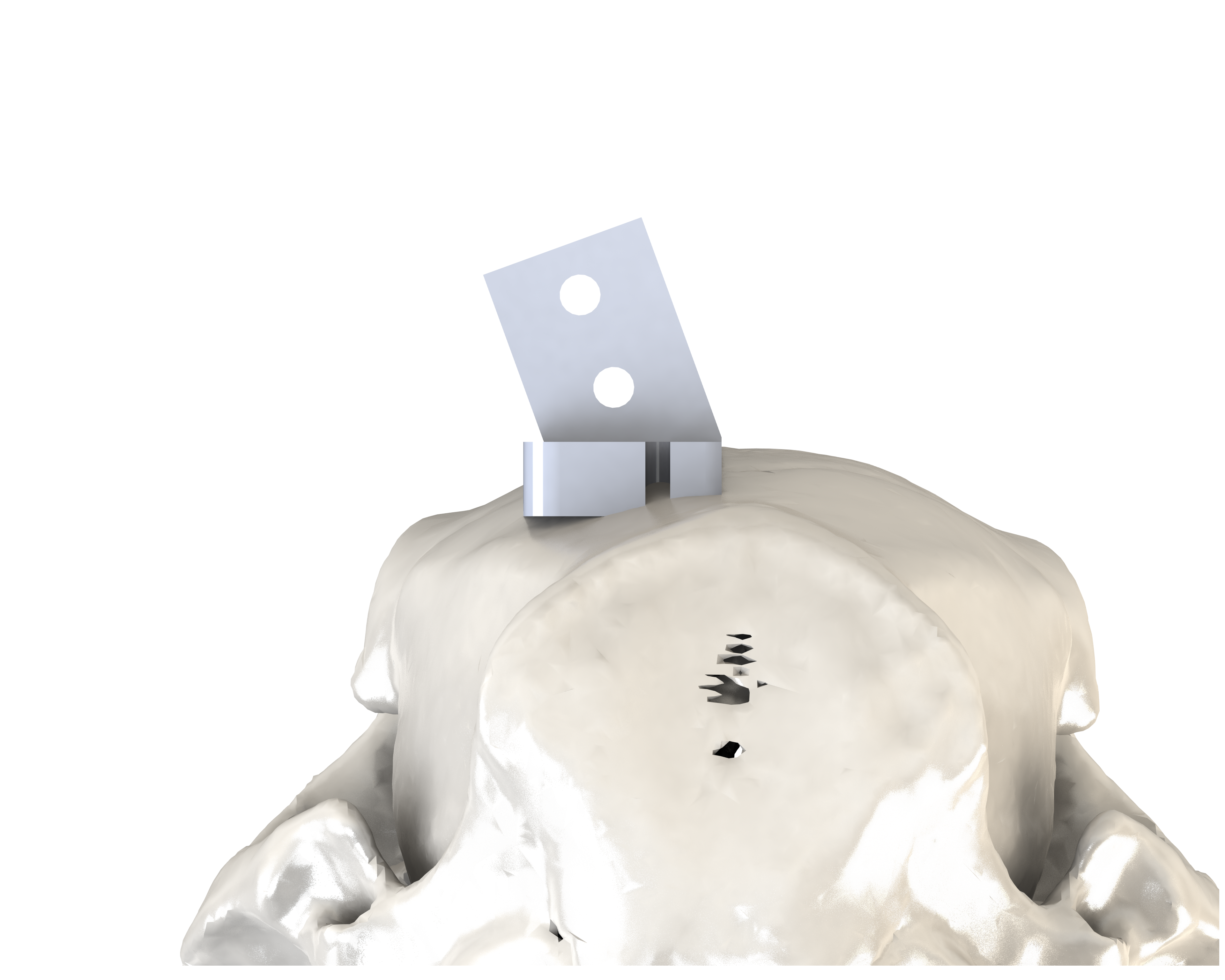

### 03_Pictures_Pig_ProbeHolder_side.png

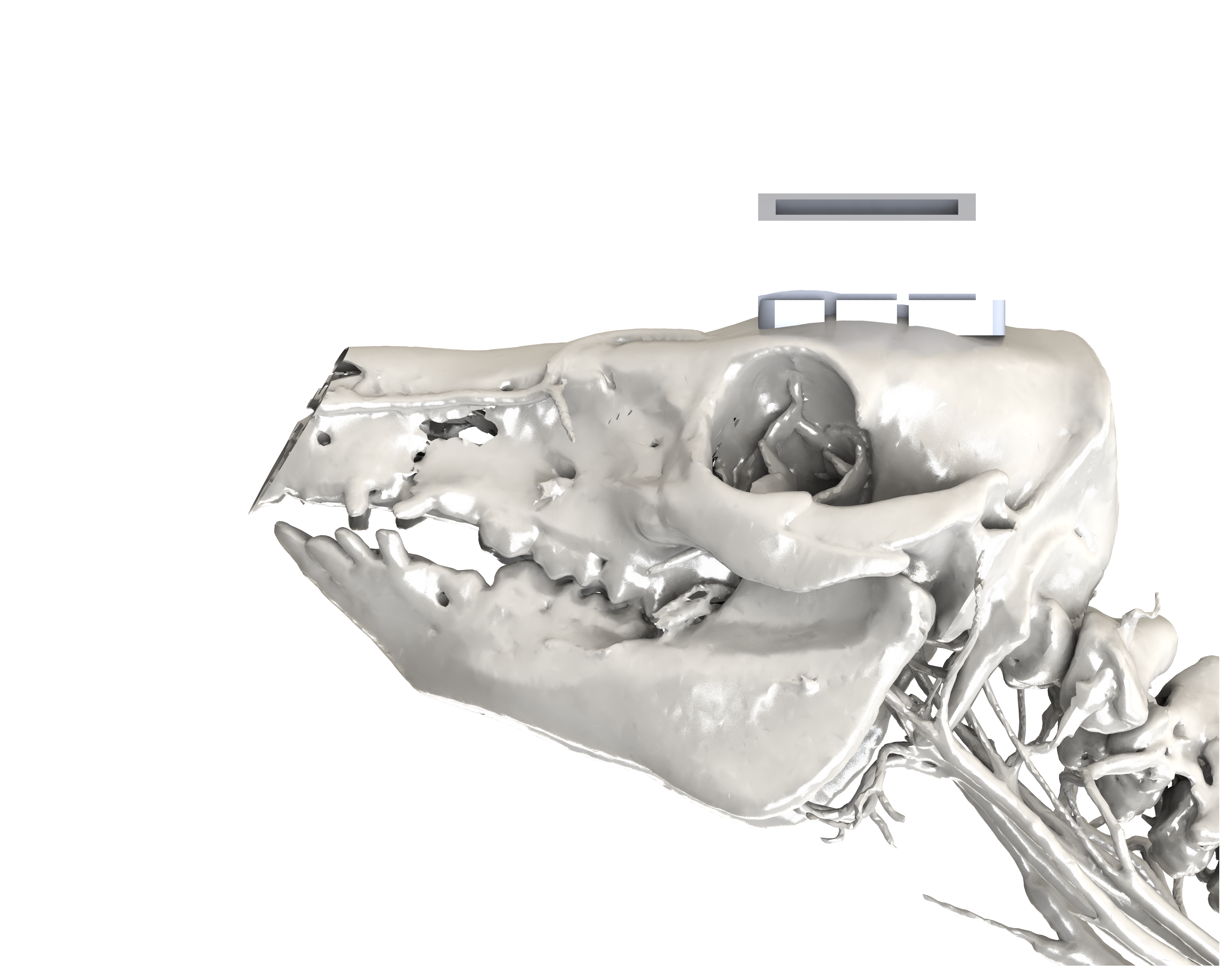

### 03_Pictures_Pig_ProbeHolder_top2.png

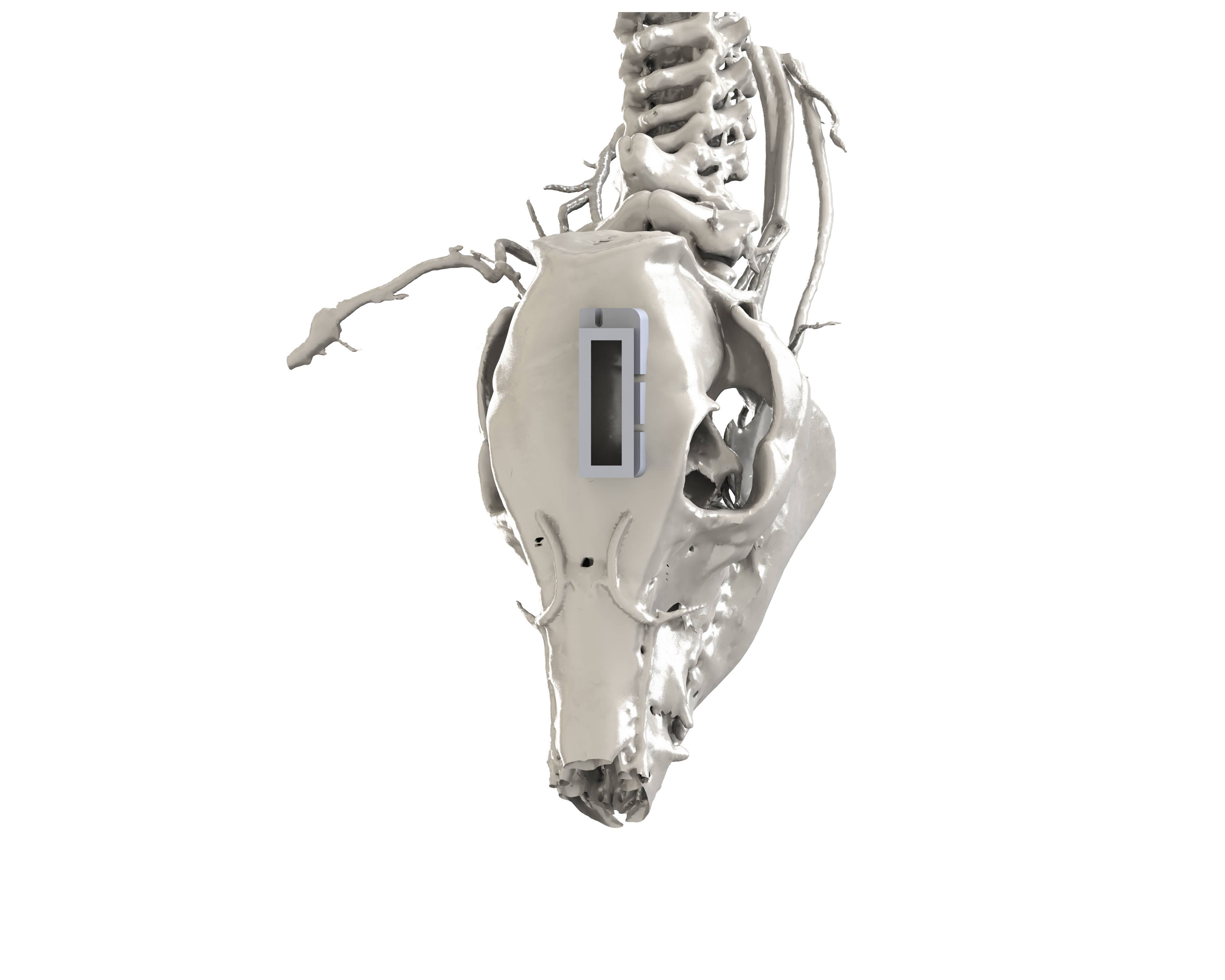

### 03_Pictures_Pig_ProbeHolder_top.png

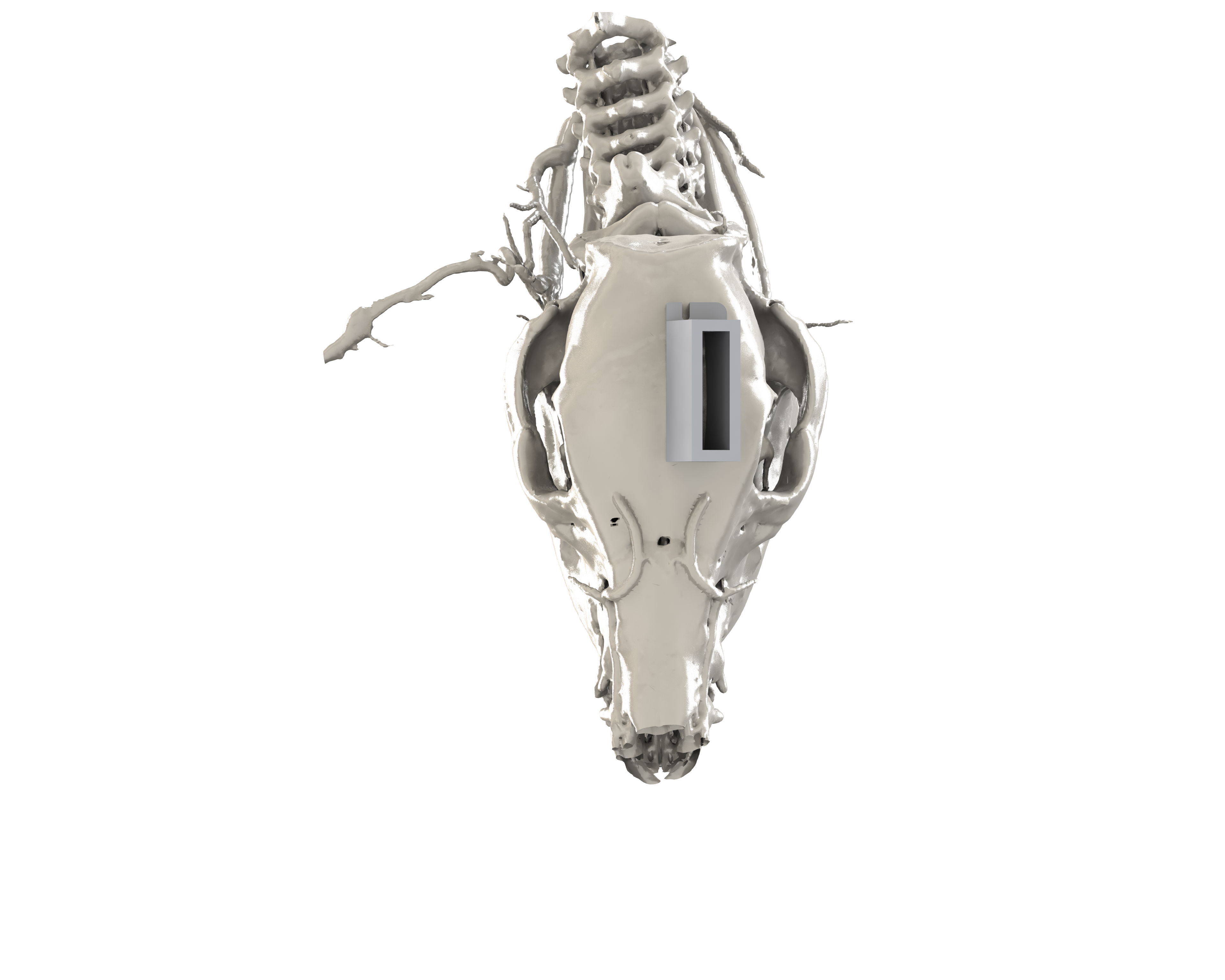

### Supplementary Figure 1

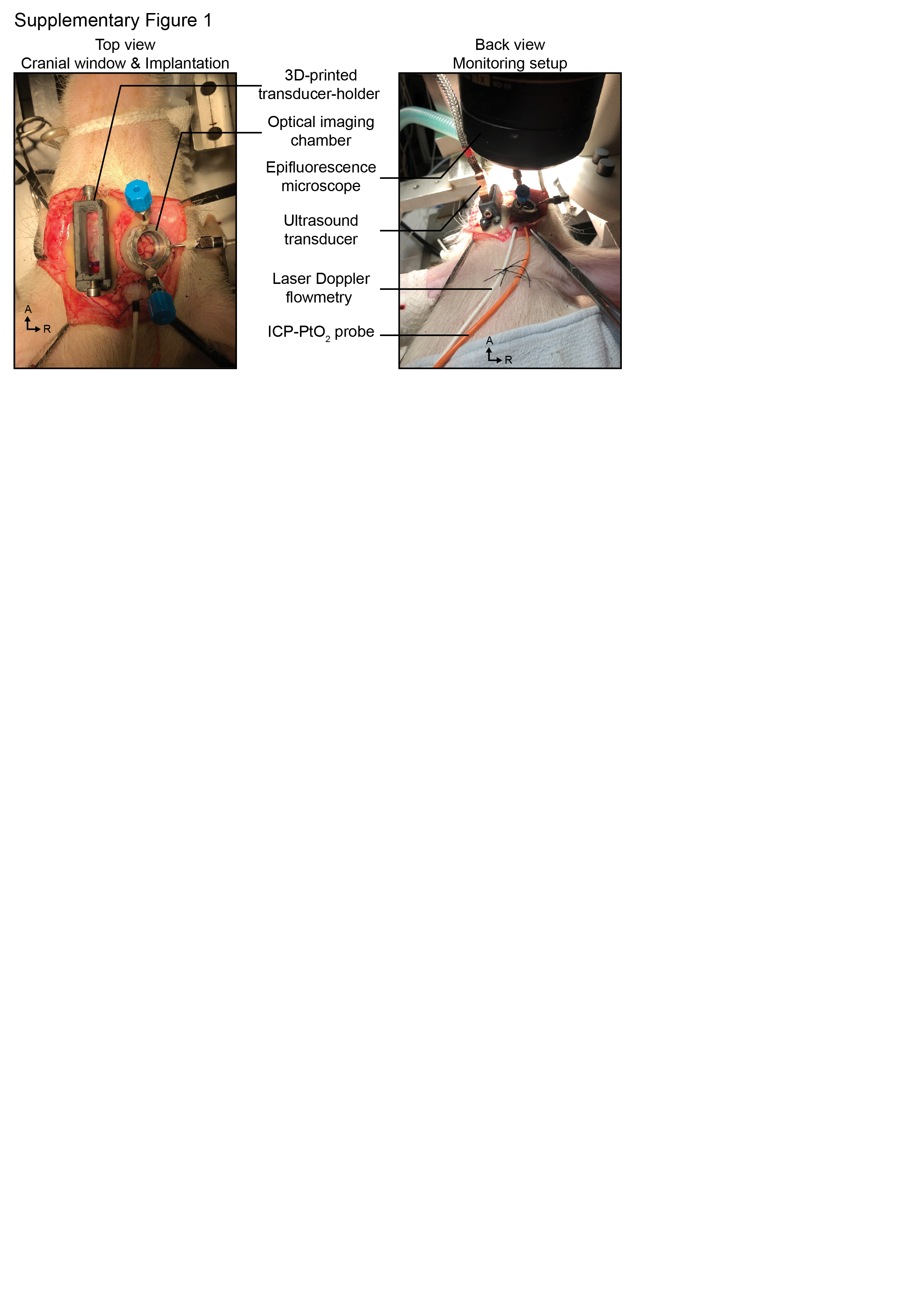

### Supplementary Figure 2

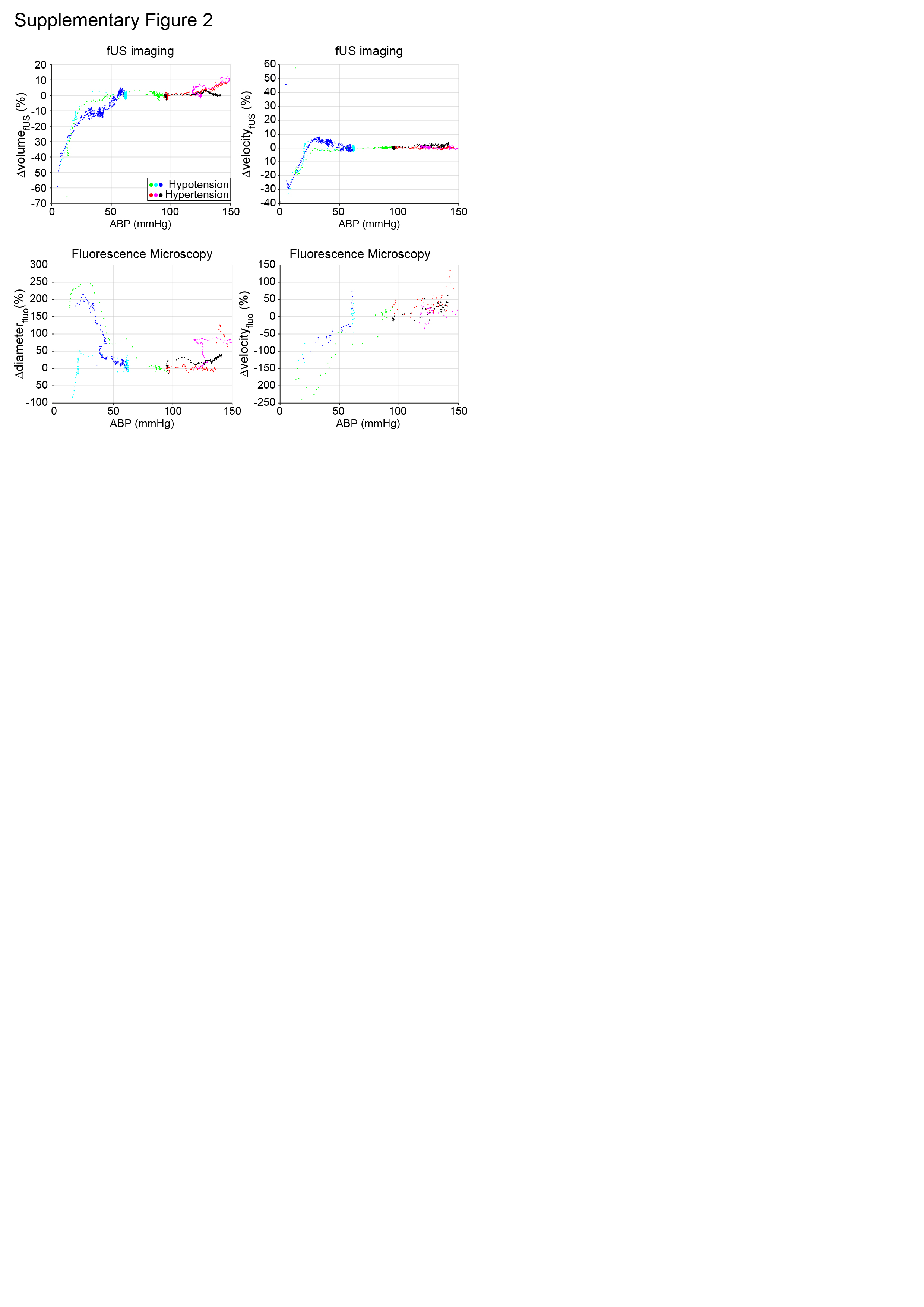

### SuppMaterial.jpg

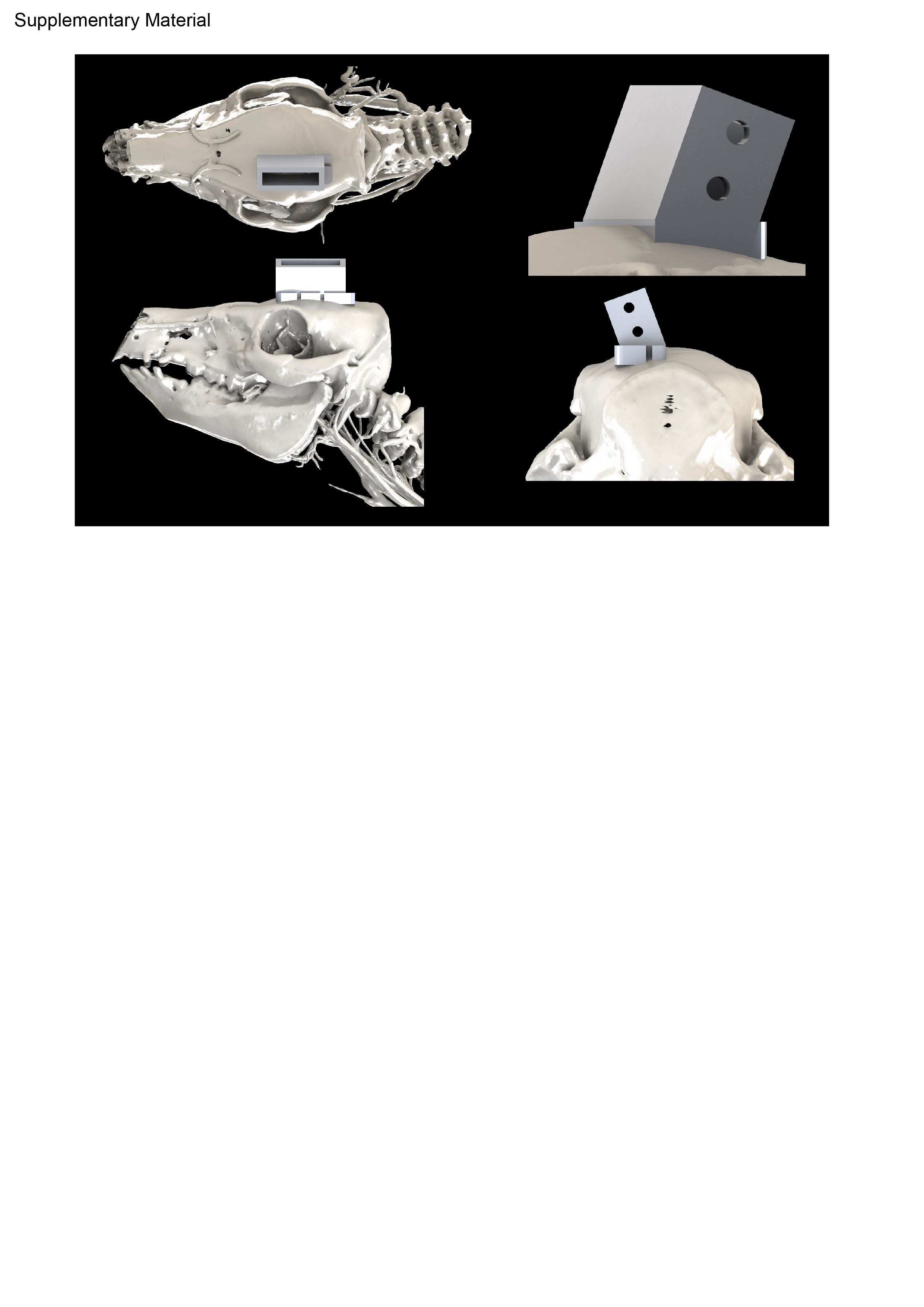
