## Supplementary material for "Continuous monitoring of cerebrovascular autoregulation using functional ultrasound imaging in the piglet brain": fUS_Sequence.docx

**Ultrasound sequence for brain monitoring.**

The ultrasound sequence used in the work has been adapted from Brunner, Grillet et al. Nature Protocol, 2021. The exact parameters and real-time processing are detailed below:

**Emission/Reception event.** During the E/R event, a plane-wave is transmitted and the backscattered echoes from the brain are received between 4 and 16µs. This time period allows echoes from 3 to 15 mm depth to be recorded. A dead time is added to give a total time of 60 µs for the E/R event. The stimulus is a 40 vpp square wave of 2 cycles.

**Plane wave acquisition.** The E/R block is repeated 3 times. The total duration of a plane-wave acquisition is 180 µs. **Processing**: The acquired data from the 3 repetitions are averaged in hardware to reduce the signal-to-noise ratio.

**Compound acquisition**. A set of 11 plane-wave acquisitions with angles (-10°, -8°, -6°, -4°, -2°, 0°, 2°, 4°, 6°, 8°, 10°) are generated in 2ms (500Hz). **Processing**: A beamforming algorithm generates a single plane-wave compound image of high resolution.

**Doppler image and spectrum.** 250 compound images are concatenated into a single Doppler image in 0.5s. **Processing**: a SVD filter is applied to the set of 250 compound images in order to remove the first 25 singular values to extract the blood signal. The spectrum and intensity are then calculated.

**Averaged Doppler image and spectrum**. The Doppler image is repeated 20 times. **Processing:** Spectrum and the intensity are averaged to reduce the signal to noise ratio.

**Brain monitoring.** The average Doppler image is repeated every 5 minutes throughout the experiment. **Processing:** At each repetition, the averaged spectrum and intensity Doppler image are stored.
